# Mapping Grip Force to Muscular Activity Towards Understanding Upper Limb Musculoskeletal Intent using a Novel Grip Strength Model

**DOI:** 10.1101/2024.12.30.630841

**Authors:** Yujun Lai, Elvera Abdel-Messih, Marc G. Carmichael, Gavin Paul

## Abstract

This work aims to evaluate a grip strength model, developed using a piecewise linear function based on the Woods and Bigland-Ritchie EMG-force model, which correlates the relationship between measured grip force and muscular activity. The grip strength model is compared against the results derived from an upper limb musculoskeletal model. Experimental results demonstrate the model’s efficacy in estimating surface electromyography (sEMG) readings from force measurements, with a mean root mean square error (RMSE) of 0.2035 and a standard deviation of 0.1207 for muscle activation (dimensionless). Moreover, incorporating sEMG readings associated with grip force does not significantly affect the optimization of muscle activation in the upper arm, as evidenced by kinematic data analysis from dynamic tasks. This validation underscores the model’s potential to enhance musculoskeletal model-based motion analysis pipelines without distorting results. Consequently, this research emphasizes the prospect of integrating external models into existing human motion analysis frameworks, presenting promising implications for physical Human-Robot Interactions (pHRI).

## 1. Introduction

The increasing ubiquity of robots in our everyday lives is a catalyst for the current shift in research towards Human-Robot Interaction (HRI), a field of research exploring the relationship between humans and robots in collaborative and cooperative scenarios [1]. The evolution of interactive systems that capitalize on human physiological signals for intent detection underscores their utility in HRI. Prior work [2] explored the inertia effects in Fitts’ law tasks within interactive environments, highlighting the nuanced ways in which human intent can manifest through muscular activity. Similarly, gamification strategies have been employed to enhance user engagement and performance in these settings [3], which provided insights into human response in interactive systems. As newer robots enable humans to safely interact with the system in close proximity, there is a focal shift towards a better understanding of the human partner’s intentions and reactions. The predominant approach to infer human intent is through the integration of human physiology in collaborative workspaces. Common methods include using physiological responses, such as bio-potentials, to deduce intent, provide feedback to the user, and hence maximize control during HRI. Commonly, this results in models that map surface electromyographic (sEMG) [4] or electroencephalographic (EEG) [5] signals to user intention and performance.

In parallel, substantial progress has been made in decoding sEMG signals to infer user intent, particularly within the domains of prosthetic control and neuromuscular function [6]–[These contributions have provided the research community with deeper insights into the neural and muscular mechanisms underlying motor control, informing both clinical applications and robotic systems. While the present study is motivated by similar challenges, it adopts a complementary approach: rather than decoding intent directly from sEMG, grip force measurements are used as a surrogate for muscular activity. This strategy enables integration with musculoskeletal modeling pipelines and has the potential to reduce dependence on dense sEMG sensor arrays, which can be impractical in dynamic human-robot interaction (HRI) scenarios.

Some frequency-domain and time-frequency-domain features have been reported to achieve good performance in detecting muscle co-contraction and predicting overall muscle strength. A part of the experimental methodology with sEMG is the variability introduced by both the users and the operators. Hudgins et al. [9] explored this effect of noise by using such frequency-domain and time-frequency-domain features to achieve a desired classification outcome for the specific actions based on sEMG signals. In their study, only one pair of electrodes was used to record sEMG signals, providing insufficient sEMG features and making it challenging to accurately pinpoint the most active muscle positions. Additionally, the electrodes were shifted in each experiment, creating variability. This underscores one current challenge: employing numerous sensors and electrodes may introduce inaccuracies over time. Although high-density sEMG (HD-EMG) systems offer rich spatial information, prior work has shown that comparable decoding performance can be achieved using a smaller number of strategically placed channels [10]. This suggests that careful sensor placement can mitigate the need for HD-EMG, which also reduces hardware complexity, setup time, and susceptibility to motion artifacts—factors that were particularly relevant for the dynamic trials conducted in this study. In addition, Bai et al. showed that rather than using high-density sEMG data, it is possible to achieve the same accuracy with as little as 5 active signal channels [10]. Hence lies the challenge for building a model for detecting muscle activity using sEMG. Such a model should be capable of identifying upper arm intent, mitigating the need for additional sensors.

### A. Musculoskeletal Models

A holistic perspective towards human physiology-driven robotics can be achieved by utilizing neuro-musculoskeletal (NMS) models. NMS models, commonly referred to as musculoskeletal models (and referred hereon as such), have long been used to simulate anatomy and physiological responses. Musculoskeletal models are created to act as analogs for biological systems and use physics-based simulations to investigate anatomical and muscular responses to input stimuli. With a variety of software available for simulations [11], [12], there is a significant body of work focusing on feature comparison [13]–[16] and real-world phenomenon validation [17]. However, there are many roadblocks in adopting musculoskeletal models in certain applications.

While simplified models can contribute towards a deeper understanding of the human central nervous system and physiological interactions, realistic analogs are hampered by the complexity of the human body. Intricate body systems, such as the upper limb complex, translate to complicated models that are challenging to validate [18]. Realistic models also pose a conundrum for researchers due to the kinematic redundancy between the number of muscles and the unconstrained Degrees of Freedom (DoF), an advantageous feature for human dexterity and adaptation [19]. With no closed-form solutions available to obtain muscular activity for a particular state, optimization-based solutions are currently the favored approach to resolving this redundancy. However, there is an ongoing debate on the appropriate cost function to be used during this optimization, with arguments for [20] and against [21] using principles of optimal control.

The most widely accepted cost function to determine muscle activation relies on the principle of minimum energy expenditure [22], deriving the minimization of the sum squared muscle activation. However, evidence suggests that this is not as relevant even for elite athletes whose successes rely on optimizing their actions [23]. Additionally, an energy expenditure-centric approach fails to consider compensatory energy usage in populations such as the elderly [24] and those with a lack of proprioceptive input such as deafferented individuals [25]. Furthermore, muscular co-contraction, a critical characteristic for human motor learning [26], is not considered when using minimum energy expenditure principles. The effect of muscular co-contractions can be seen in the regulation of body segment impedance [27], accuracy [28], and task variability [29], which highlights current limitations.

A possible solution to tackle current limitations is integrating novel information available through emergent technologies. These aim to supplement or supplant parameters and functionalities of musculoskeletal models using extrinsic channels of information, such as camera-based 2D & 3D human pose estimation [30], marker-less motion capture systems [31], and IMU-based motion analysis [32].

Following the same line of thought, models that bridge the gap between task-centric measures and physiological states can complement musculoskeletal models. Such models incorporate some form of physical contact between the human and the robot, especially for physical HRI (pHRI) applications.

A handle is commonly used to facilitate end-effector-based interactions [33]. Besides the interaction forces at the end-effector, the grip strength on the handle can be used as a channel of information, similar to kinematic velocities during unconstrained voluntary point-to-point movements [34]. More recently, their potential to tackle human-centric robotics is realized with works leveraging the model’s results for robotic assistance [35], [36].

With multiple confounding factors affecting human motor control [37], learning [38], and planning [39], musculoskeletal models have been invaluable for researchers to isolate and investigate their influence on human response. The fidelity of musculoskeletal models in representing human muscular activity is pivotal for their application in predictive models of human behavior. Recent work by Sutjipto et al. [40] introduces a novel representation of strength profiles, offering a more granular understanding of the relationship between grip strength and muscle activation. This advancement is crucial for refining musculoskeletal models to better predict human physiological responses in dynamic tasks.

### B. Human Grip Strength

Human grip strength has been well studied with excellent inter-rater and test-retest reliability [41]. Early studies have shown its links to the somatosensory system [25], with pressure distribution dynamics investigated by Gurram et al. [42]. Despite being a tertiary source of information, grip strength has the advantage of overcoming biases associated with sEMG readings, especially with dynamic movements, which are particularly challenging for sEMG-based models. Additionally, they can be computationally accounted for as defined by the musculoskeletal models.

The use of grip strength provides the opportunity for a model that maps the relationship between grip strength and muscle activation for the relevant muscle groups. Finger flexion and extension are attributed to the *flexor digitorum profundis* (FDP) and *extensor digitorum communis* (EDC) muscle groups [43], with the FDP muscle group being the primary contributor to human grip force.

In pHRI applications, grip strength has been used to indicate human physiological response to perturbations [45]. Still, the inherent variability among human demographics creates a challenge for intent detection and human-robot role arbitration [46]. Human performance can vary based on psychological and physiological characteristics, and thus, alternative perspectives on inferring intent are necessary to provide robust solutions.

One popular model hypothesized for the relationship between muscular activity (% normalized sEMG) and force generation (% MVC) is the two-element EMG-Force model by Woods and Bigland-Ritchie [44] (Fig. 1), which suggests that there are intrinsically two types of muscle fibers during muscle activation (see Fig. 1). This model is supported empirically [47] while De Luca et al. identified that the curve parameters are unique to each muscle group [48]. Assuming that the slow Type 1 fibers generate force asymptotically at 30% of MVC, the Woods and Bigland-Ritchie model can be parameterized using piecewise equations to isolate the two fiber types.

**Fig. 1:**
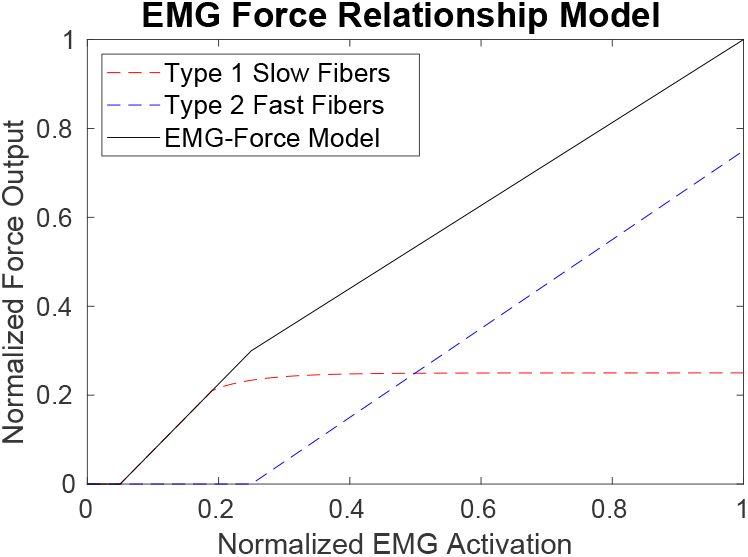
The two-element EMG-Force model proposed by Woods and Bigland-Ritchie used for the grip strength model (reproduced from [44]).

This work aims to build a grip strength model based on the Woods and Bigland-Ritchie model to correlate the relationship between grip strength and muscular activity. Such a model may alleviate the need for extraneous sensors to be attached to the human user, allowing for seamless integration of extrinsic information channels into the human motion analysis pipeline. Experimental data will validate the grip strength model, using measured grip force and sEMG signals as analogs of grip strength and muscular activity. The influence of the grip strength model on a typical human motion analysis is investigated through a commonly used musculoskeletal model [49], [50].

The rest of the article is as follows: Section II outlines the experiment methodology to create and validate the grip strength model, with results presented in Section III. A discussion on the implications of the results follows in Section IV, followed by the conclusion in Section V.

## II. Experiment Methods

Ten male adults, aged between 18 and 30, and presenting no neuromuscular disorders, participated in the experiment. The experiment protocols were explained to the participants before obtaining their informed consent. This research has been carried out according to the ethical guidelines of the Human Research Ethics Committee at the University of Technology Sydney (UTS HREC approval no.: ETH18-3029).

The experimental method was composed of three types of trials: a reference trial, isometric trials and exercise trials. The reference trial was used to determine the MVC of each participant, the isometric trials were used to construct the grip strength model, and the exercise trials were used to validate the model. The isometric trials were randomised between 33%, 66% and 100% to and the original aim was to affirm MVC - but this was not possible when the data was sorted.

Statistical and analytical methods used in this study include a one-way ANOVA to evaluate the presence of muscle fatigue over time, stepwise linear regression to fit the grip strength model, and RMSE as the primary error metric for model validation. All analyses were conducted using MATLAB (Math-Works Inc., USA), including functions such as *pwelch, fft*, and *stepwiselm*. Grip force and sEMG data were normalized using reference trial maxima to enable inter-subject comparison. A series of double-blind, randomized trials were conducted to obtain each participant’s isometric and dynamic grip force across three voluntary contraction levels: 33%, 66%, and 100%. A set of standardized verbal instructions was provided to participants, with participants instructed to maintain 90^*°*^ elbow flexion, no radial/ulnar deviation of the wrist, and a natural grasp on the handheld dynamometer. Across all participants, the estimated wrist extension was within *±* 15^*°*^ with no significant changes throughout the experiment. While maximal grip strength is typically associated with a wrist posture closer to 30° extension [51], a neutral posture was adopted here to simplify setup and reduce participant fatigue across repeated trials.

Participants rested their forearm on a table to maintain a consistent and repeatable posture without mechanical fixation of limb segments. This configuration allowed for simplified setup and signal acquisition, removing the need for real-time joint tracking or inverse kinematics during isometric trials. While this introduces a potential source of variability, it enabled a focus on sEMG and grip force signals. Postural consistency was supported through verbal instruction and table-based support, ensuring approximate maintenance of the intended joint configuration throughout the task.

The experiment consisted of one reference trial, 12 isometric trials, and 6 exercise trials. In each trial, the grip force exerted by the participant was measured by a digital dynamometer (*±* 0.1*kg*). The measurements from the digital dynamometer were captured using video footage and extracted manually post-hoc. Administrators advised participants against reading their grip force results after each trial, although no physical restrictions were employed to enforce this. For all trials, the maximum grip force measurement was collected. This aligns with the sEMG data processing pipeline (Section II-E) to extract the maximum Root Mean Square (RMS) values.

In the reference trial, participants were instructed to “squeeze as hard as possible for 5 seconds while maintaining your trunk and arm posture”. The sEMG readings from this reference trial were set as the MVC muscle activation for the rest of the experiment. They were further instructed to “use this [*reference trial*] to relate to the 33% and 66% grip strength level for the rest of the experiment”. The participants approximated the 33% and 66% grip strength levels to the best of their ability. No feedback on muscular activity was provided to the participants during the experiment. The only feedback provided was when a trial had been completed.

### A. Participant Measurements and Sensor Placement

Prior to the recorded trials, the length of each participant’s ulna and humerus were measured using identical protocols as those from the CDC (US) [52] and the NHS (UK) [53].

Muscular bio-potentials are collected through silver-silver electrodes using the Delsys Bagnoli 8-Channel system (Delsys, Natick, MA). Electrodes are positioned based on SENIAM guidelines^1^, targeting muscle groups listed in Table I.

**TABLE I:**
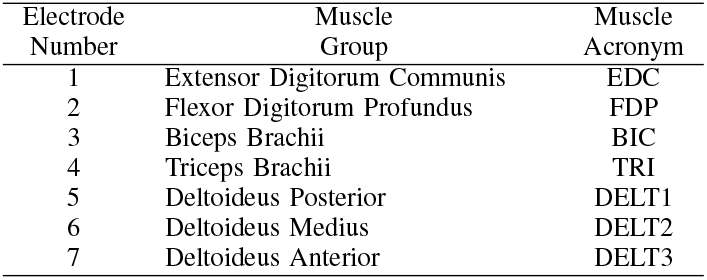
The targeted muscle groups for the placement of the sEMG electrodes.

Motion capture software was utilized to track the kinematics of each participant’s upper limb. A total of 12 Optitrack Prime cameras, in conjunction with the Motive software (Natural-Point, Corvallis, USA), tracked four user-defined rigid body assets (Fig. 4). Each rigid body asset consists of 3 passive reflective markers with known locations to provide unique poses. The rigid body assets are placed to coincide with the 4 major body segments of the model: chest (thorax), upper arm (humerus), forearm (ulna), and hand (metacarpals).

### B. Isometric Trials

Participants were requested to maintain the arm and trunk posture for isometric trials as shown in Fig. 4. In each trial, participants were given a 5-second visual countdown, after which participants were instructed via visual instructions to squeeze the dynamometer for 5 seconds at 3 distinct strength levels: 33%, 66%, and 100%. Between each trial, participants were given a one-minute break to mitigate muscular fatigue arising from the experiment.

Each participant performed 4 trials per grip strength level, with the order of the 12 trials randomized using Latin squares. The grip strength level for each trial was occluded from the administrator using a screen not visible to the administrator. For each isometric trial, sEMG data collection commenced 2 seconds into the countdown, resulting in 8 seconds of data for each trial.

### C. Exercise Trials

At the start of the exercise trials, participants were instructed to watch a video that shows a reaching exercise^2^. The exercise trials were performed in a darkened room to reduce reflections from the passive reflective markers. Thus, LED light strips were diffused and attached around the LCD display of the digital dynamometer to facilitate the reading of the grip force measurements on the video footage (see Fig. 2(a)).

**Fig. 2:**
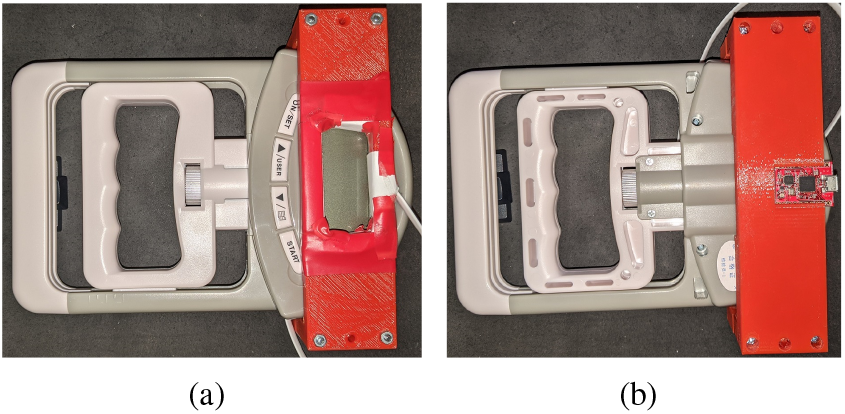
The front and back views of the digital dynamometer fitted with a 3D printed part to house (a) the diffused LED light strip, and (b) the attitude and heading reference system (myAHRS+).

Similar to isometric trials, visual instructions for the grip strength levels were given to participants. They were verbally instructed to “squeeze at the indicated grip strength level and start your [*sic*] movements while maintaining your [*sic*] trunk posture”. No further requirements, such as task completion speed or accuracy, were imposed on participants.

Kinematic data for participants are collected from the exercise trials using the rigid body assets, starting when the countdown commences. Since there were no task completion speed requirements imposed, participants were instructed to indicate to the administrator verbally when they “feel like you [*sic*] have completed the exercise from the video”. Similar to the isometric trials, sEMG data collection commences 2 seconds into the countdown. Both sEMG and kinematic data collection are stopped manually by the administrator when they receive participants’ verbal indication.

Both kinematic and sEMG data from the exercise trials are used for post-hoc analysis with the musculoskeletal model. External forces exerted on the participant’s hand are calculated from linear acceleration readings from an attitude and heading reference system (WITHROBOT, Seoul, Korea), along with the mass of the sensor-equipped dynamometer (*m* = 0.516*kg*). These forces are applied to the musculoskeletal model at the lunate coordinate of the hand segment. Forces induced by gravity are also included in the dynamic analysis of the musculoskeletal model (Section II-G).

### D. Incorporation of Muscle Fatigue Effects

Participants were provided with a minute break between each trial. To ensure muscle fatigue does not confound the results, a Power Spectral Density (PSD) plot was generated for each set of trials. Any shift in the peak power frequency over the experiment indicates long-term muscle fatigue, which may skew the results. The raw sEMG signals from the experiment were analyzed using both Fast Fourier Transforms and Welch PSD estimate, functionalities performed using MATLAB’s implementation (*fft* and *pwelch* functions, respectively). Gaussian smoothing with a window size of 40 was performed on the Welch PSD to obtain clear peaks of the PSD curve. For each trial, the sEMG signals were segmented into three temporal sections, with the first section starting 0.5s after the visual instruction was provided to the participants.

The PSD estimates for the first and last two 100% strength level trials (across both isometric and exercise trials) were used to perform a one-way ANOVA analysis to determine the effects of longer-term muscle fatigue.

### E. sEMG and Kinematic Data Processing

The collected sEMG signals are rectified before applying a 4th-order Butterworth lowpass filter with a cutoff frequency of 4Hz. The first and last 0.5 seconds of the processed sEMG signals are ignored to remove artifacts arising from observations of partial phases. An overview of the sEMG processing pipeline can be seen in Fig. 3.

**Fig. 3:**
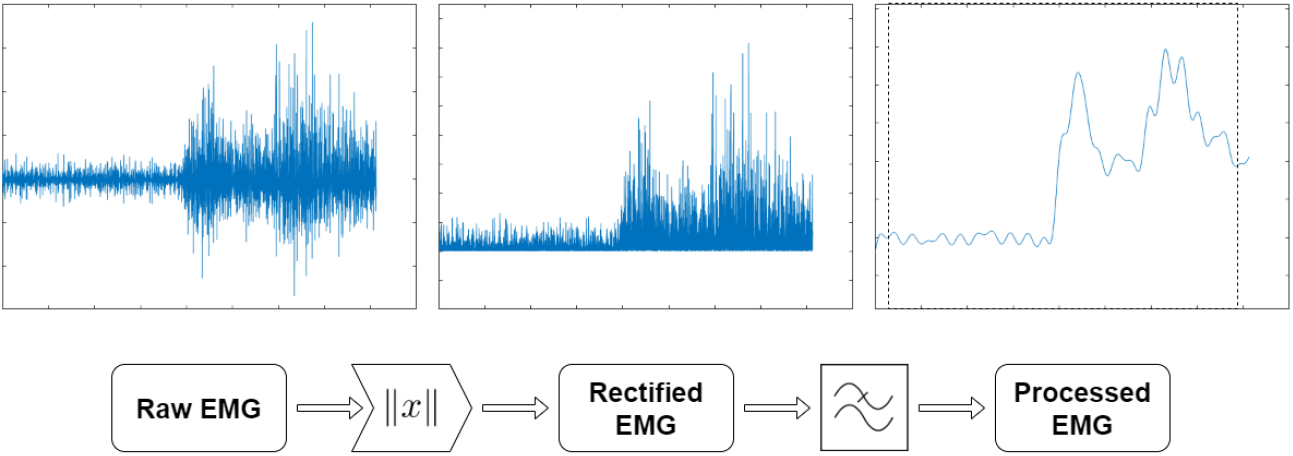
An illustration of the sEMG processing pipeline.

**Fig. 4:**
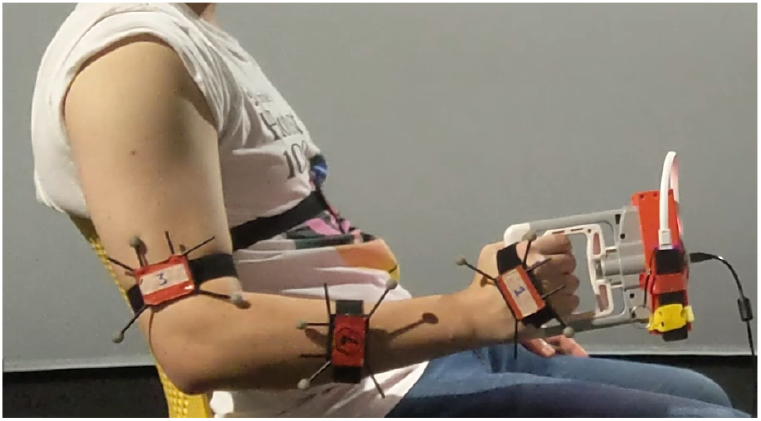
The posture participants were requested to maintain during isometric trials. Four rigid body assets were attached to the arm (3) and torso (1) as a reference for the musculoskeletal model.

The grip strength model aims to build a mapping between muscular activity and the grip force exerted by each participant. The sEMG signals are normalized to allow for quantitative analysis between participants. A common method to normalize sEMG signals is to conduct Maximum Voluntary Contraction (MVC) trials, which aim to maximize the muscle activation of specific muscle groups. Despite ongoing work to determine the optimal set of exercises for inter-rater reliability and MVC values [54], different exercises are prescribed for different muscle groups.

### F. Grip Strength Model

To construct the grip strength model mapping the relationship between grip strength output and muscular activity, isometric trial data from all participants are used to generate and validate the model through exercise trials. To ensure the model’s validity across all participants, both sEMG intensity and grip force were *range-normalized* by dividing each trial’s value by the maximum value recorded during the participant’s reference trial (i.e., their assumed MVC). This normalization allows inter-participant comparisons by scaling data to each individual’s maximum effort.

The validation of the grip strength model is performed using dynamic movements, which can introduce variability in the results. To assess the effect of changing posture during the exercise trials, a comparison of geometric muscle arms was performed across three distinct poses: the starting pose, the point at which the hand is furthest from the torso, and an intermediate in-transit pose. The mean absolute difference in moment arms was calculated for 41 muscles across 94 musculoskeletal model coordinates, including those associated with the FDP and EDC muscle groups. These coordinates represent local reference frames for insertion points and bones; joints provide the degrees of freedom (DoFs) and are treated as distinct entities in OpenSim models. Although other posture-dependent muscle parameters — such as the force–length–velocity (FLV) relationship and pennation angle — also influence torque, these were not explicitly modeled in this analysis.

In essence, the isometric trials were used to build the model, and the exercise trials were used to validate the model.

### G. Musculoskeletal Model

The upper limb musculoskeletal model used for the experiments is derived from the Stanford VA Upper Limb model [49] with updated model parameters from live human data [50]. While more recent models [55] include finer digit-level actuation and passive joint properties, our study focuses on grip strength as a scalar measure of intent involving the major flexor and extensor groups (FDP and EDC), which remain common to both models. The model consists of 15 degrees of freedom and 50 Hill-type muscle-tendon units (MTUs) [56]. Modifications were made to the model to simplify the model while maintaining bodies, coordinates, and actuators relevant to the experiment. Updates to the model include^3^:

- Inertial properties of the humerus, radius, and capitate were set using anthropomorphic estimation for the average male height and weight [57]. Principal moments of inertia for the body segments are taken from [58].
- The MTU model of the original model [59], is replaced by the more recent [60] MTU model. Available MTU parameters are transcribed to the new MTU model (maximum isometric force, optimal muscle-tendon length, pennation angle at optimal length, tendon slack length). Other MTU parameters (Force-Length-Velocity curves) are set to the default values in OpenSim.
- The model complexity is reduced by restricting motion to 5 DoFs, comprising of: shoulder flexion/extension, abduction/adduction, internal/external rotation (3 DoF), elbow flexion/extension (1 DoF), and wrist flexion/extension (1 DoF). The remaining joints are locked to match the experiment posture (e.g., closed digits, neutral pronation/supination, and 0^*°*^ wrist abduction/adduction). MTUs that do not contribute to the 5 DoFs are disabled by ignoring them during calculations. MTUs that do not contribute to the 5 DoFs are disabled by ignoring them during calculations.
- Only the MTUs contributing directly to grip strength—specifically those associated with the FDP and EDC muscle groups—were retained for analysis. This decision is supported by the model’s moment arm matrix, which indicates that other muscle groups have minimal influence on grip force in the postural configuration studied. This simplification is aligned with the pHRI context of this study, where grip strength serves as a scalar and interpretable proxy for user intent.

For each participant, the measurements of their humerus and ulna are used to scale the geometry of the model manually. The length of the humerus and ulna matches their respective measurements, while the length of the radius is proportionally scaled based on the ulna measurement.

For each exercise trial, external forces exerted by the dynamometer were calculated by utilising the linear accelerations and applied at the metacarpals. The upper limb motion analysis is conducted by performing inverse kinematics, inverse dynamics, and kinematic analyses. All three operations utilize the functionalities in OpenSim.

The generalized forces obtained from inverse dynamics are used for static optimization in MATLAB (using the *fmincon* function) to obtain an estimate of the muscular activity, 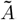,during the exercise trials. A minimum sum of squared activation cost function is used:

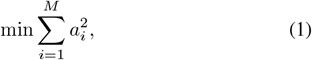

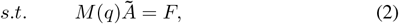

where *M* (*q*) is the computed moment arm matrix, which depends on the current kinematic state, *F* is the net generalized forces on the model joints, and 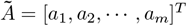 is the activation level (0 → 1) for the *m* MTUs in the model.

For the purpose of demonstrating the feasibility of integrating extrinsic information channels with musculoskeletal models, the muscle activations for FDP and EDC are assumed to be identical (assuming isometric co-contraction). The muscular activity estimate is obtained using the normalized sEMG readings of the FDP muscle group. The same sEMG readings are applied to multiple MTUs in the musculoskeletal model, as we assume sEMG readings are derived from the muscle groups. This consolidation of the muscle group is outlined in Table II. The generalized forces caused by the EDC and FDP muscle groups, as calculated from the measures sEMG readings, are removed from the optimization process, that is, *F*_*opt*_ = *F* − *F*_*sEMG*_. The optimization is then performed using the modified moment force matrix. The modifications remove the MTUs contribution from the EDC and FDP muscle groups from the optimization.

**TABLE II:** The replacement of the MTU activation in the musculoskeletal model with the model estimate.

| Muscle Group | Model MTUs |
| --- | --- |
| EDC | EDCI, EDCL, EDCR, EDCM |
| FDP | FDPI, FDPL, FDP R, FDP M |

## III. Results

From an initial analysis of the reference trials, the common method to normalize the sEMG readings was not followed as 6 out of the 10 participants demonstrated supramaximal sEMG readings during the isometric and exercise trials for electrodes 1 & 2. This is likely due to the lack of feedback provided to participants throughout the experiment.

Thus, a different normalization method is used based on the sEMG intensity range. Since participants were instructed to hold onto the handheld dynamometer (0.516kg) in a relaxed manner, the sEMG intensity for muscle activity during this time is assumed to be the minimum value.

The sEMG intensity is obtained by partitioning the signal into rolling windows which are 0.25s long (0.05s apart), and calculating the RMS for each window. Since the experiment includes an exercise suitable for MVC (e.g., 100% grip strength), each participant’s normalization range is formed by the minimum RMS value between 0.5-3.0s and the maximum RMS value across all trials.

The sEMG signals from electrode numbers 3–7 were not processed since they are used for qualitative analysis with the static optimization results.

### A. Contribution of Muscle Groups

The generation of human grip strength is attributed to the muscle groups that control finger flexion and extension: FDP and EDC. To ascertain the influence of each muscle group on a user-generated grip force, a series of one-way ANOVA was conducted between each muscle group and their relationship to the measured grip force.

Only the subset of isometric trials where participants were instructed to exert at 100% grip strength was used in this particular analysis, with the muscular activity posited to be a better representation of grip strength. Isometric trials for 33% and 66% grip strength are assumed to incur influence on muscular activity as participants try to control and maintain the instructed grip strength level. This can be seen in Fig. 5, where there is a larger spread of muscular activation despite a smaller grip force spread.

**Fig. 5:**
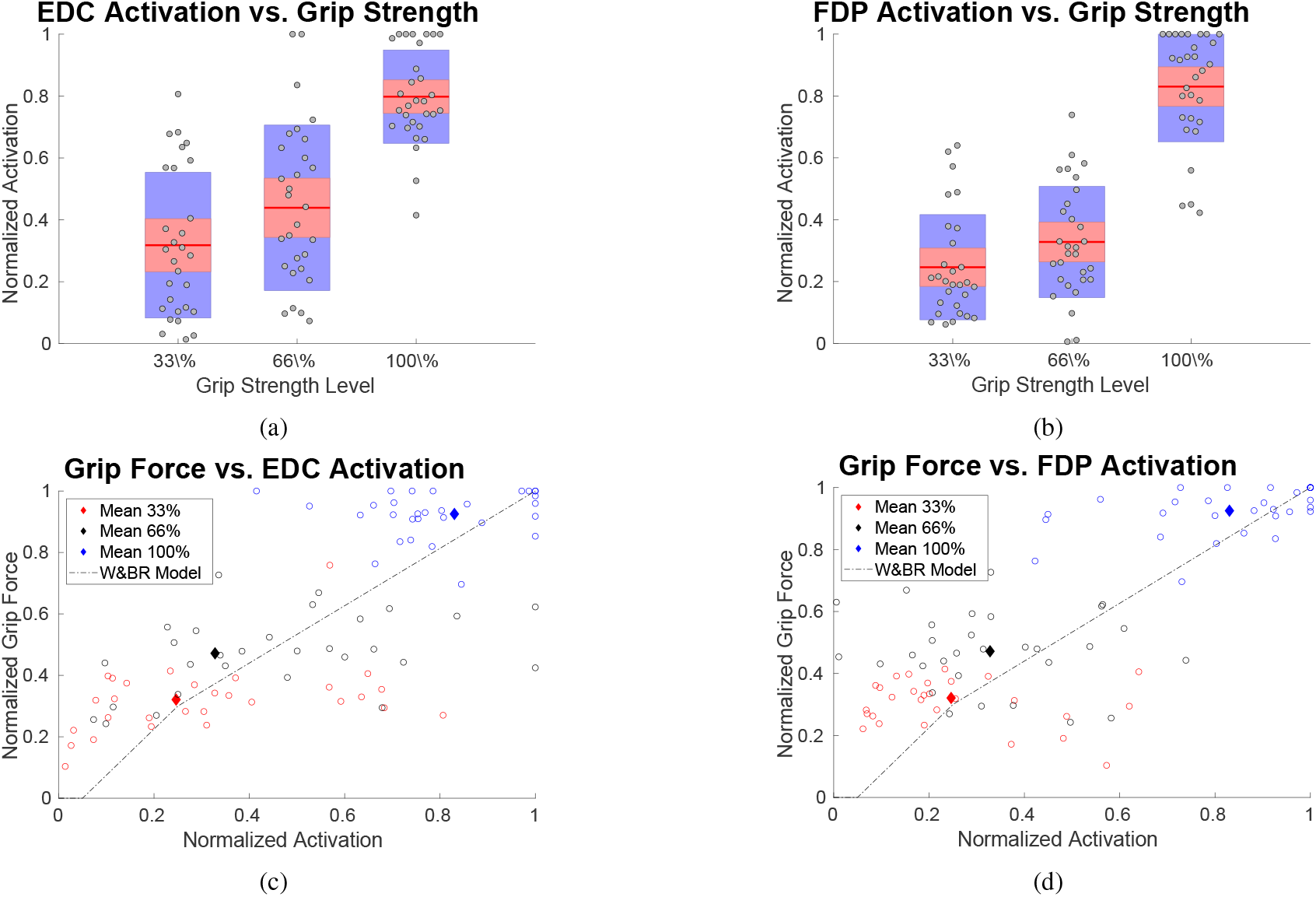
The relationship between the grip force measurements and muscle activation for all participants in the experiment. The Woods and Bigland-Ritchie muscle EMG-force model is given as a reference. Each dot represents a single isometric trial, with four trials per grip strength level (33%, 66%, 100%) per participant. Data from 10 male participants are shown (a total of 120 trials). Each participant’s individual results and identity are not visually distinguishable.

The normalized sEMG activation of FDP and EDC are statistically the same (*F* (1, 54) = 1.0570, *p* = 0.3085), with no significant difference during 100% isometric trials. This is reinforced by the Pearson and Spearman’s Rank Order correlation score, which shows a relatively equal correlation between each muscle group and the measured grip force as outlined in Table III.

**TABLE III:** Correlation scores between EDC, FDP, and grip force. Pearson’s correlation scores are in the top right triangle, while the lower left triangle shows Spearman’s Rank Order correlation score.

|  |  | EDC Muscle | FDP Muscle | Grip Force |
| --- | --- | --- | --- | --- |
| | | Pearson's $\rho$ | | |
| EDC Muscle | Spearman's $\rho$ | | 0.60 | 0.73 |
| FDP Muscle |  | 0.56 |  | 0.77 |
| Grip Force |  | 0.72 | 0.68 |  |

Given the comparable correlation between measured grip force and the FDP and EDC muscle groups, the decision was made to adopt a single-variable model. The preference for utilizing the FDP muscle group is attributed to its broader sEMG signal range during contractions, facilitated by range normalization of the signals. Using a higher range mitigates the effects of signal noise, alleviates the chances of supramaximal and negative sEMG activation, and accommodates a diverse participant pool with varying muscular strengths.

### B. Grip Strength Model Validation

Two participants were excluded from further analysis due to consistently flat sEMG signals across all trials, indicating a likely electrode placement issue during setup. These signals lacked observable muscle activity and showed no correlation with grip force.

Visual inspection of the remaining isometric trial data (Fig. 6) identified two additional participants whose data deviated systematically from the expected trend. Their normalized sEMG activations exhibited a negligible or negative correlation with grip force, preventing reliable model fitting. These exclusions were based on full-trial inconsistencies rather than isolated outlier points.

**Fig. 6:**
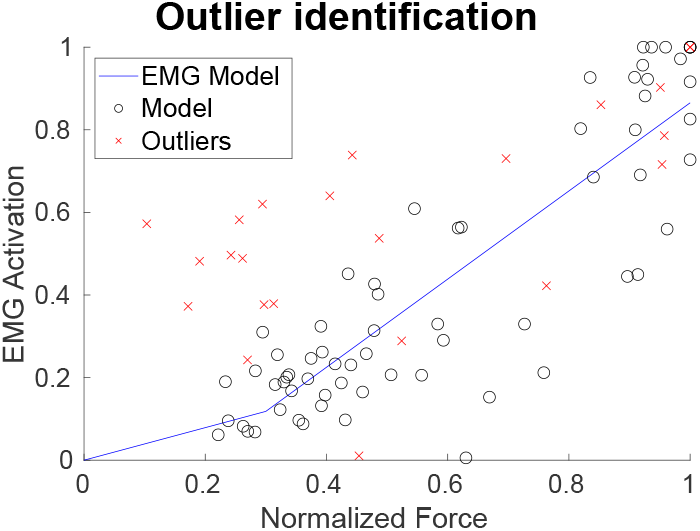
A scatter plot showing the relationship between normalized grip force and FDP activation across all isometric trials. Black circles indicate the trials retained for model fitting, while red crosses denote outlier trials that were excluded. Outliers include trials from two participants with flat or inconsistent sEMG responses across all conditions, resulting in low or negative correlation. The fitted EMG-force model is shown in blue.

For the remaining participants, the relationship between grip force and FDP activation was modeled using a step-wise linear regression approach (*stepwiselm* function in MATLAB). Trials from the excluded participants produced root mean square errors (RMSEs) exceeding three standard deviations above the mean. The final grip strength model, built using the remaining data, achieved a mean RMSE of *µ* = 0.1512 with a standard deviation of *σ* = 0.1207.

The grip strength model was validated using the results from the exercise trials (Fig. 7), showing a mean RMSE of *µ* = 0.2035 with a standard deviation *σ* = 0.1207. While evidence suggests a power curve would fit a similar relationship [42], this was not investigated in this work. A comparison between the instructed grip strength level and the measured grip force is shown in Fig. 8. While the 33% and 100% levels show good alignment with expected values, the 66% condition is consistently underestimated across participants. This likely reflects the difficulty of estimating submaximal effort without feedback, as participants received no guidance during the trials. Rather than indicating an experimental error, this deviation highlights the variability in the subjective perception of effort, particularly at intermediate levels.

**Fig. 7:**
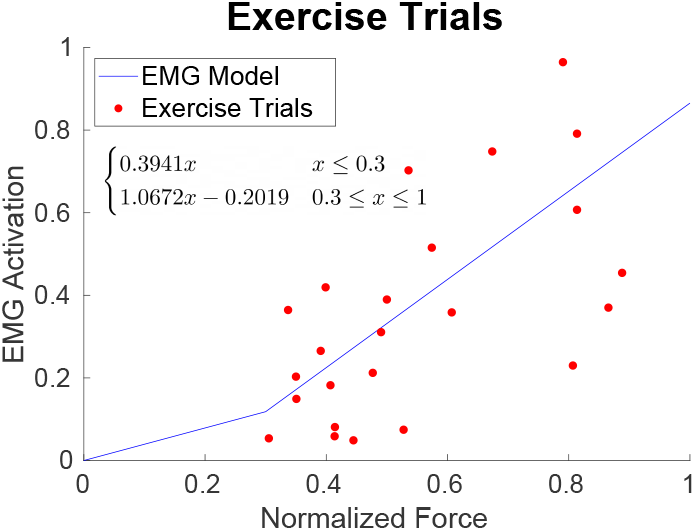
Sample data from valid exercise trials and the parameterized grip strength model used for validation.

**Fig. 8:**
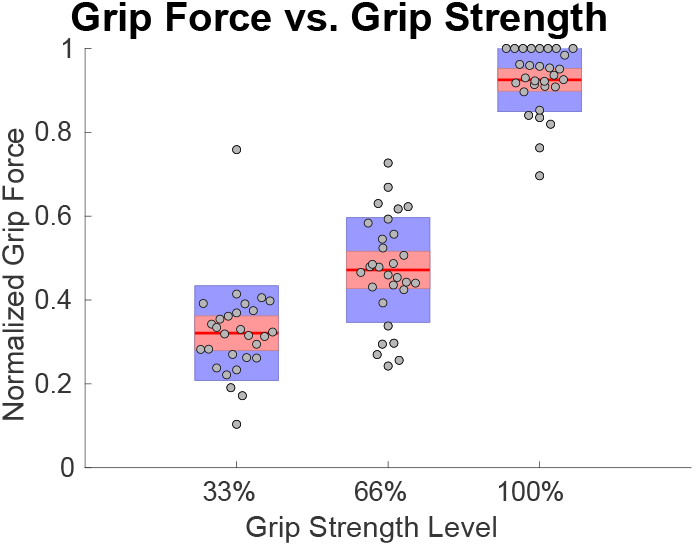
A modified box plot of the instructed grip strength level and the measured grip force for all participants. The red line indicates the mean, the standard deviation is shaded in pink, and the 95% confidence interval is shaded in blue. The discrepancy at the 66% level reflects the participants’ subjective estimation without feedback, which introduced variability around this submaximal target.

For the three poses evaluated, the mean absolute differences in moment force between the starting pose, and the ending and in-transit poses are *µ*_*end*_ = 1.6913*N* (3.76% of the total) and *µ*_*transit*_ = 1.2963*N* (2.82% of the total), respectively. This showed that participants’ moment force differences had a minor effect during the exercise trials.

### C. Impact of sEMG Data Inclusion on Muscle Activity Estimates in Grip Strength Modeling

A series of optimizations using MATLAB’s *fmincon* function were conducted to investigate the influence of the grip strength model’s estimates on resultant muscle activity. The cost function for the optimization follows the default used in OpenSim, the minimum sum squared value of the muscle activation. Due to the use of minimization during the optimization process, the resultant muscle activity is generally underestimated when compared to empirical data. The discrepancies seen in Fig. 9 for the FDP and EDC muscle groups can be attributed to the range normalization process for the sEMG readings and implicit errors in the process of data collection. Moreover, simplified external load estimates from the dynamometer may not fully account for participant compensatory behaviors.

**Fig. 9:**
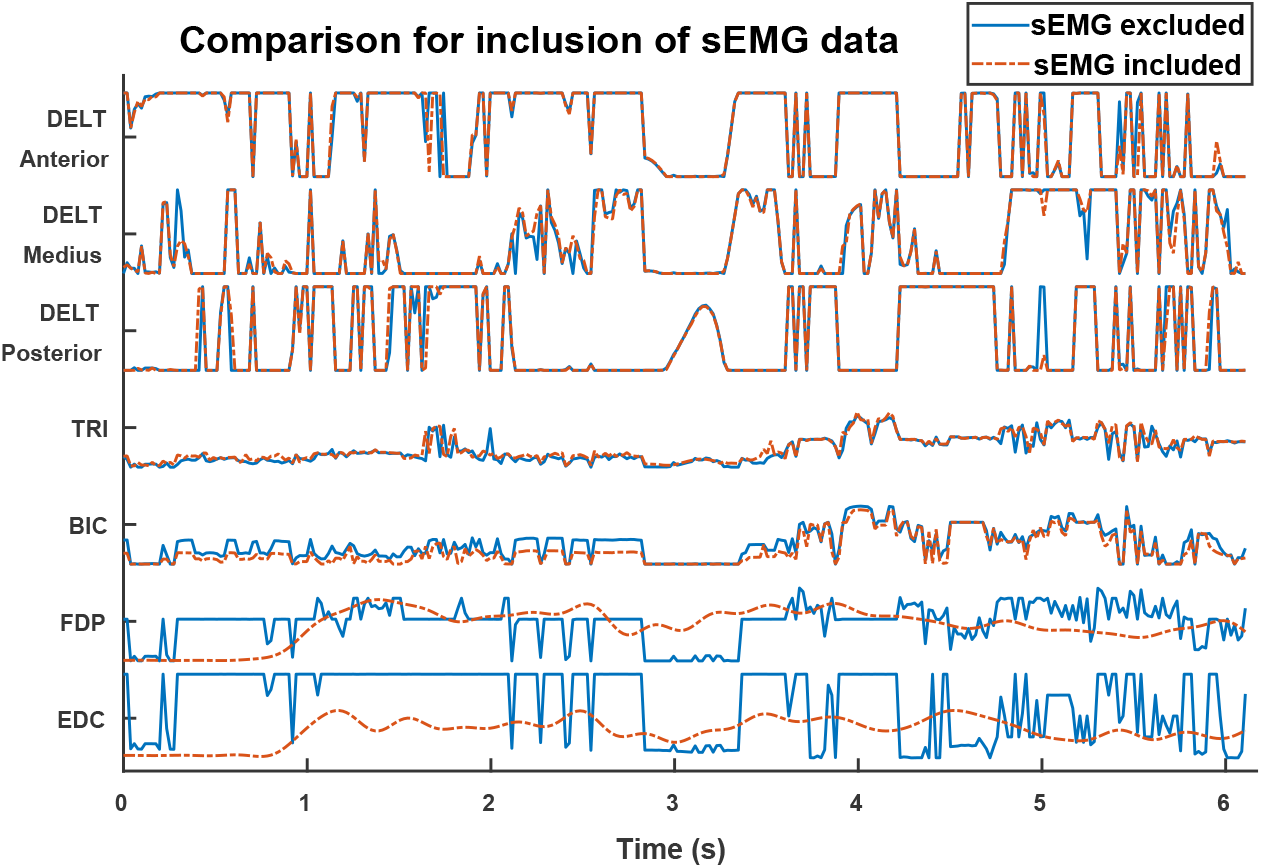
A comparison of the effect of the grip strength model estimate on the muscle activation levels derived from static optimization. The traces shown are outputs from the musculoskeletal model, not raw sEMG signals. The saturation observed reflects the optimization outcome, not measurement clipping. Only the FDP and EDC activations (orange) are derived from sEMG input; the others (e.g., biceps, triceps, deltoids) are optimized estimates. This comparison highlights how incorporating sEMG-derived estimates affects proximal muscle activations in the model.

Figure 9 provides a visual contrast between scenarios with sEMG data inclusion and those without, particularly noticeable in muscle groups such as FDP and EDC. This distinction underscores the significant role that sEMG data plays in refining the accuracy of muscle activity predictions. Notably, the variations in muscle activity over time, as illustrated, offer a nuanced understanding of the dynamic interplay between different muscle groups during grip strength exertions.

The inclusion of sEMG data significantly enhances the model’s sensitivity to muscle activation patterns, particularly within the FDP and EDC muscle groups, as evident in the pronounced activity profiles displayed in the figure. This enhancement demonstrates the value of sEMG data in capturing the complex dynamics of muscle activity during grip tasks, despite the challenges of range normalization and potential measurement errors. The observed discrepancies between empirical data and model estimates highlight the important role of accurate sEMG integration in improving muscle activity modeling.

### D. Assessment of Muscle Fatigue Effects

The PSD plots across the three temporal sections for all participants indicated that acute effects of muscle fatigue are present within each trial (Fig. 10). This phenomenon is expected as participants are instructed to exert grip force for at least 5 seconds in each trial. Thus, to assess the longer-term effects of muscle fatigue, only the first temporal section of each trial was used for analysis.

**Fig. 10:**
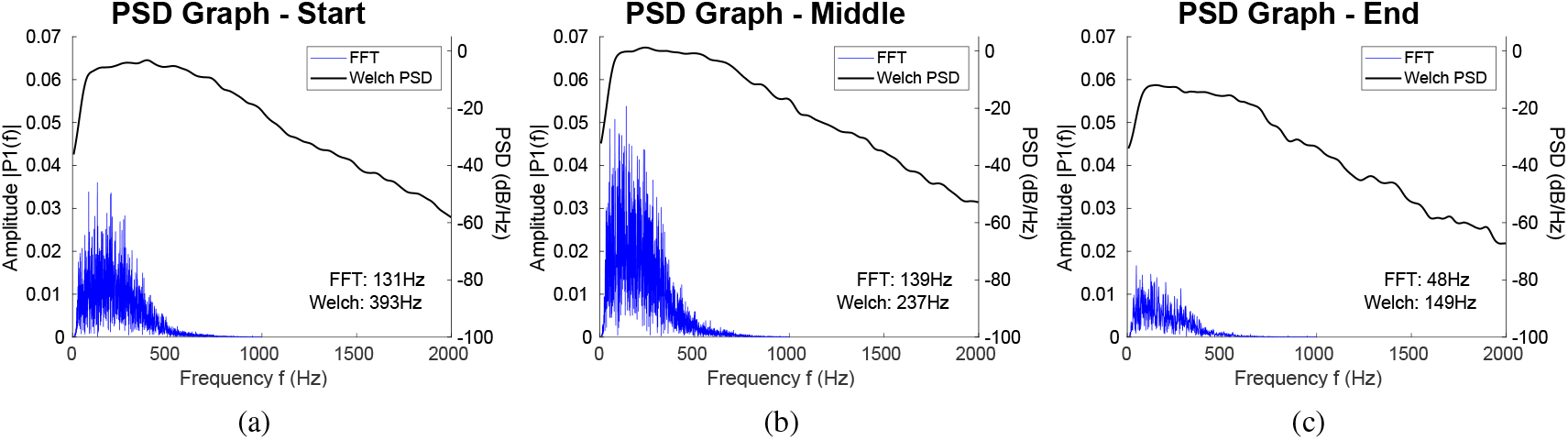
The Power Spectral Density (PSD) plots across (a) the first third, (b) the second third, and (c) the final third of a trial. These plots are representative of the PSD across most of the isometric and exercise trials.

Across all participants, the one-way ANOVA analysis indicates that there was no significant shift in the peak frequency (FFT: *F* (1, 38) = 0.9943, *p* = 0.3250, Welch: *F* (1, 38) *<* 0.001, *p* = 0.9823), indicating that the effects of muscle fatigue are negligible in the experiment.

## I.V Discussion

From the experiment’s sample size, the generated grip strength model has proven to be a feasible adjunct for estimating sEMG activation based on measured grip force. Despite utilizing data from only 10 participants, the correlation between each muscle group and the measured grip force aligns with the Woods and Bigland-Ritchie model, as illustrated in Fig. 5. The current understanding of Force-Length-Velocity (FLV) effects on MTUs is well-documented [61]. While this study does not explicitly model the combined effects of posture-dependent muscle force and moment arms, we acknowledge that both factors are important contributors to joint torque and should be considered together in future analyses. Empirical visual observations further corroborate that the Woods and Bigland-Ritchie model can effectively link sEMG activation with grip force. In our study, this model served as a conceptual reference for developing the empirically derived grip strength model, but it was not implemented as a computational constraint in the musculoskeletal simulation, which instead relies on the Millard equilibrium muscle model. The wrist posture was held approximately neutral across participants, although prior studies suggest that maximal grip strength typically occurs closer to 30° of wrist extension [51]. This study did not control or vary wrist angle, and future iterations of the model could incorporate this biomechanical factor to further refine predictive accuracy.

Visual inspection of isometric trial data (Fig. 6) revealed outliers, suggesting potential experiment setup deficiencies. An initial analysis focused on the impact of these outliers, indicating an enhancement in model quality upon excluding certain data points. However, this study encountered a limitation due to data loss during retrieval. For some trials, the video footage of the experiment was underexposed, leading to an inability to read the measured grip force (∼ 14%) from the video footage. Despite this, the analysis of the remaining dataset revealed a positive correlation between grip strength levels and normalized grip force. This empirical correlation between grip force and muscle activity is consistent with prior findings in the literature, such as Mogk and Keir [62], who demonstrated that forearm muscle activity during gripping can be predicted using grip force and joint posture in regression models. These findings affirm the model’s validity, extending the Woods and Bigland-Ritchie model to represent sEMG activation to muscle fiber force output. The model can be extended to infer human intent, limiting the use of additional sensors—marking a novel contribution to the field.

Visual observations of Fig. 7 and Fig. 8 reinforce the relationship between the instructed grip strength and the actual grip force measured during the exercise trials. The data collected in this study primarily covers normalized grip forces above 0.3, which limits the conclusions that can be drawn about low-force behavior. While this may exclude fine motor tasks or activities of daily living, the model is intended for physical human-robot interaction (pHRI) contexts, where grip force serves as a stabilizing or communicative signal rather than for precision control. The consistent and predictable trends observed among participants suggest that the model remains valid for moderate to high grip force interactions and could be used in conjunction with other models for broader applicability. Furthermore, this study used a single, fixed-handle design typical of constrained pHRI scenarios. While this limits generalizability to tasks involving variable object sizes, it allows for better repeatability and is representative of many practical HRI configurations.

The model was developed and validated using data from male participants only, consistent with the demographic basis of the musculoskeletal models employed. Further work is needed to investigate the model’s applicability across broader populations, including female subjects or individuals with different anthropometric profiles.

Although a set of electrodes and sEMG sensors were initially employed, the model’s results indicate alternative sensors to sEMG systems can be utilised which are more practicable in pHRI applications. This highlights the model’s capability to supplant traditional sensor-based measurements. This advancement simplifies experimental setups and opens new research methodologies where reliance on sEMG sensors can be reduced or eliminated. Further, this enhances efficiency and broadens the scope for future muscle activity measurement and modeling explorations.

The result in Fig. 9 indicated that including the grip strength model has not affected the functionality of the motion analysis pipeline for a musculoskeletal model for the muscles in the proximal sections of the upper limb. This supports the concept of employing a suite of discrete models that provide extrinsic channels of information to supplement musculoskeletal models during motion analysis. A notable phenomenon is that muscle activation from FDP and EDC may alleviate the activation necessary from biceps brachii to enable the recorded motion. This reinforces the previous observation of bias that may be introduced into musculoskeletal models by supplementary information sources and needs to be explored in future works considering the applications of this model.

Considering the insights from the experimental results, where discrepancies in muscle activity estimation were observed due to the dynamometer’s limitations and other factors, a potential improvement could be introduced. Implementing a lightweight force transducer in place of the dynamometer could mitigate the identified effect, enhancing the accuracy of the grip strength model’s estimates and reducing the observed discrepancies in muscle activity. This adjustment suggests a pathway to refining the experimental setup for more precise and reliable outcomes.

The grip strength model aims to contribute to the current trend in generating supplementary information channels that complement musculoskeletal models. This will encourage their adoption in emerging fields such as rehabilitation [37], assessment [63], and pHRI [30]. While the grip strength model provides additional information sources when inferring human intent, such as in pHRI, it may introduce bias in the muscular activity analysis. Realistic musculoskeletal models are particularly sensitive to this possibility as the upper limb complex has complicated muscle-tendon paths.

Building on the foundation laid by this study, future endeavors could focus on refining and personalizing the grip strength model to cater more effectively to individual physiological variations. The prospect of tailoring the model using advanced computational techniques, such as machine learning algorithms or Bayesian statistics, presents an exciting frontier. This personalized approach would enhance the model’s accuracy and applicability in diverse scenarios and open new pathways for integrating real-time physiological responses into comprehensive human motion analysis frameworks. Such advancements could improve the precision of assessments in rehabilitation and physical human-robot interactions, ensuring that the grip strength model evolves into an even more potent tool for predicting and understanding muscle activity.

## V. Conclusion

This work has successfully developed and validated a grip strength model that maps the intricate relationship between measured grip force and muscular activity. Through testing with 10 participants across a range of isometric and dynamic gripping tasks, the presented model has demonstrated accuracy in estimating sEMG readings from grip force measurements, with a mean RMSE of 0.2035 and a standard deviation of 0.1207. The findings validate the model’s effectiveness in integrating with upper limb musculoskeletal analyses, confirming that the inclusion of sEMG data does not compromise the optimization of muscle activation or the overall outcomes of motion analysis. This achievement underscores the model’s robustness and potential to augment musculoskeletal modeling by minimising sensory equipment.

Moreover, the application of the presented grip strength model in practical scenarios highlights its value in enhancing the precision of muscle activity estimations, particularly in the domain of pHRI. The model’s capacity to infer upper muscle intent based on grip force alone, aligning with established sEMG-force models, represents a significant stride toward more intuitive and accessible musculoskeletal analyses. Through examination and interpretation of experimental data, the assessment of muscle activity patterns, this work confirms the model’s reliability and strategic relevance in the broader context of human motion studies. This approach paves the way for more sophisticated analyses that can contribute meaningfully to advancements in rehabilitation, assessment, and pHRI, affirming the grip strength model as a critical tool in the intersection of biomechanics and interactive technologies.

## Footnotes

1 http://www.seniam.org/

2 The exercise video shown to participants is available here

3 Updated model: https://github.com/y-lai/modifiedUpperLimbModel.

